# Divergent marine anaerobic ciliates harbor closely related *Methanocorpusculum* endosymbionts

**DOI:** 10.1101/2024.03.12.584670

**Authors:** Anna Schrecengost, Johana Rotterová, Kateřina Poláková, Ivan Čepička, Roxanne Beinart

**Author notes:** Corresponding authors (AS); (RB). Department of Marine Sciences, University of Puerto Rico Mayagüez, Mayagüez, Puerto Rico, USA. **Competing Interests:** The authors declare no competing financial interests.

## Abstract

Ciliates are a diverse group of protists known for their ability to establish various partnerships and thrive in a wide variety of oxygen-depleted environments. Most anaerobic ciliates harbor methanogens, one of the few known archaea living intracellularly. These methanogens increase the metabolic efficiency of host fermentation via syntrophic use of host end-product in methanogenesis. Despite the ubiquity of these symbioses in anoxic habitats, patterns of symbiont specificity and fidelity are not well known. We surveyed two unrelated, commonly found groups of anaerobic ciliates, the Plagiopylea and Metopida, isolated from anoxic marine sediments. We sequenced host 18S rRNA and symbiont 16S rRNA marker genes as well as the symbiont ITS region from our cultured ciliates to identify hosts and their associated methanogenic symbionts. We found that marine ciliates from both of these co-occurring, divergent groups harbor closely related yet distinct intracellular archaea within the *Methanocorpusculum* genus. The symbionts appear to be stable at the host species level, but at higher taxonomic levels, there is evidence that symbiont replacements have occurred. Gaining insight into this unique association will deepen our understanding of the complex transmission modes of marine microbial symbionts, and the mutualistic microbial interactions occurring across domains of life.

---

Microbial eukaryotes, or protists, are among the most common and abundant organisms on the planet, representing most of the diversity of eukaryotic life. Many protist lineages have evolved to live without oxygen, and these anaerobic protists frequently form symbioses with bacteria and archaea (1,2). For those with a fermentative metabolism, symbioses with hydrogen-scavengers may have facilitated transitions to an anaerobic lifestyle (1,3). In particular, many anaerobic protists host methanogens in their cytoplasm which are often found adjacent to host mitochondria-related organelles (MROs). These MROs generate energy for the host via fermentation, ultimately producing H2 which the symbionts utilize as a substrate for methanogenesis. This syntrophic transfer is thought to reduce intracellular hydrogen tension, increasing host metabolic efficiency and enabling relatively large, highly active grazers to thrive in anoxia (4–6). Despite their ubiquity and diversity in oxygen-depleted habitats ranging from marine sediments to gastrointestinal tracts, basic information about these partnerships, such as the factors that influence which bacterial or archaeal lineages they partner with, is poorly known.

Ciliates are particularly well-suited for life without oxygen: virtually all ciliate lineages include representatives adapted to full or partial anaerobic lifestyles, and most of the recent research on free-living anaerobic protists centers around ciliates (3,4,7–9). However, little is known about symbiont diversity and specificity within and among host ciliate lineages, especially those from marine environments. Most studies have focused on a few host species isolated from a handful of geographically isolated, mainly freshwater, habitats (3). Even so, there is evidence that these associations are stable and may be vertically transmitted: the symbionts divide synchronously in one host species (*Plagiopyla frontata*) (10), and endosymbiotic methanogens persist in host resting cysts (11). Additionally, identical symbiont species have been detected in a single ciliate species isolated from geographically distant locations (7,12). On the other hand, the close phylogenetic relationships observed between methanogenic endosymbionts and their free-living relatives suggest that endosymbiont replacements have occurred during their evolution (8,9,11). Importantly, studies to date have shown no evidence of co-diversification which would be typical of vertically transmitted symbionts (11,13). At the same time, interpretation of these studies is confounded by many variables, including a diversity of geographic locations, habitats, and host phylogeny.

Here, we surveyed the symbionts of two obligately anaerobic groups, the Metopida (Armophorea, SAL) and Plagiopylea (CON3P), which are common in oxygen-depleted marine habitats. These two groups are phylogenetically divergent (14) but co-occur in the shallow, intertidal sediments we sampled here (Table S1, Figure S1). Therefore, we minimized geographical and environmental differences, allowing us to focus on differences in symbiont specificity both within and among these two groups. We identified the methanogenic symbionts from 31 different *Plagiopyla* and 14 different *Metopus* isolates, representing 13 species-level ciliate lineages (Figures 1, 2, S1; Supplemental Information) (15). We found that, despite their large phylogenetic diversity and geographic range, all of these ciliates harbor closely-related *Methanocorpusculum* symbionts whose 16S rRNA gene sequences are 98-100% identical (mean 99%∓0.54 s.d.), with one exception (Figure 1). *Methanocorpusculum* has primarily been recorded in wastewater environments and as an endobiont of animals and anaerobic ciliates (7,12,16,17). Transmission electron microscopy (TEM) revealed that the *Methanocorpusculum* symbionts from representatives of both host groups are highly integrated with the host cell, located in between host MROs (Figure 1). These findings are similar to those from studies of *Methanocorpusculum* in other ciliates, suggesting a tight integration with the host on both a structural (13) and genomic (7) level is a feature which distinguishes this genus from other symbiotic methanogens.

**Figure 1.**
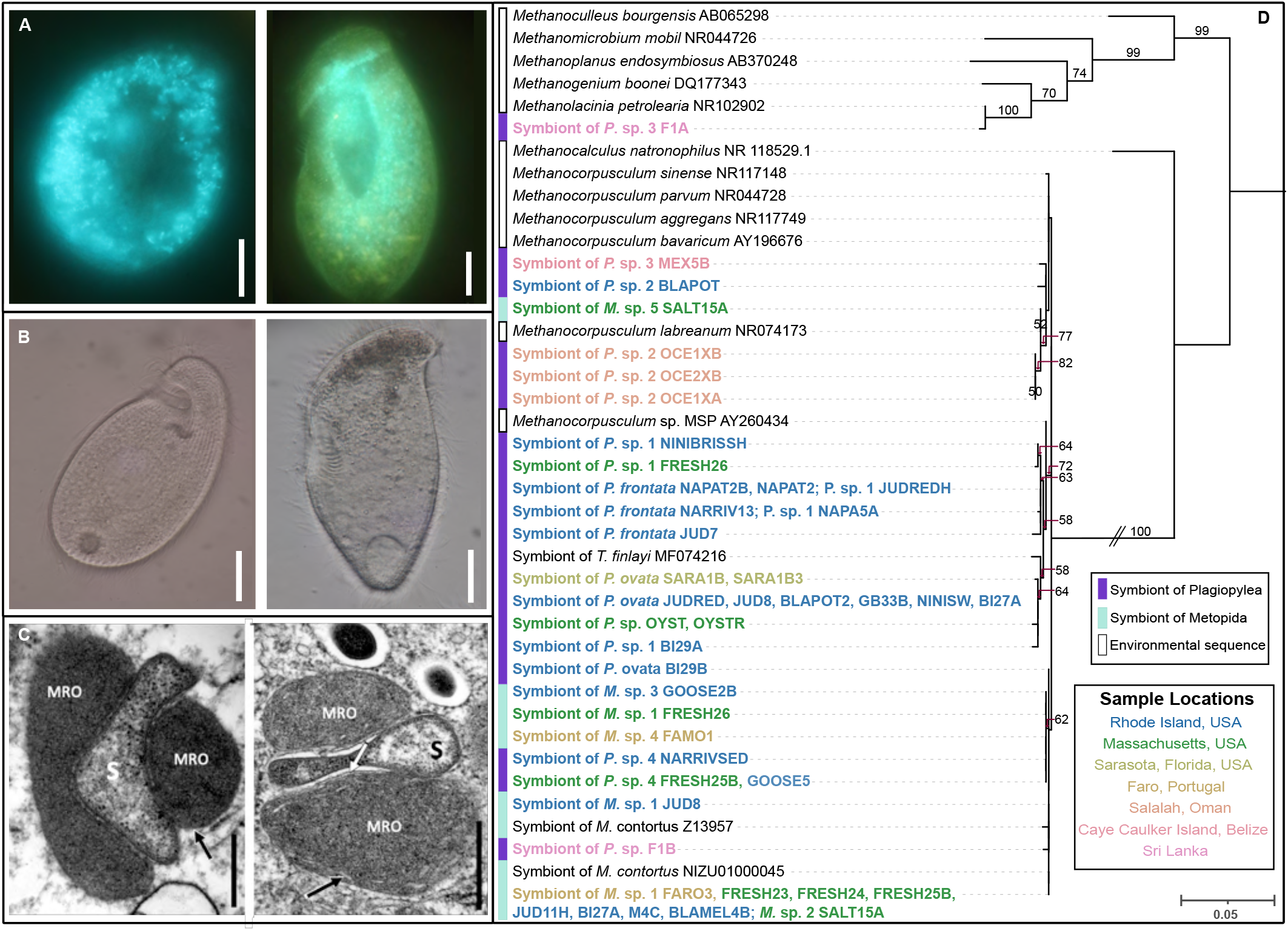
(A) Micrographs depicting autofluorescence of methanogenic symbionts within host *Plagiopyla* (OYST, left) and *Metopus* (FRESH26, right) cells. (B) Representative light micrographs of host *Plagiopyla* (NAPAT2, left) and *Metopus* (BI27A, right) cells. (C) Transmission electron microscopy (TEM) depicting the ultrastructure of methanogenic symbionts within host *Plagiopyla* (FRESH26, left) and *Metopus* (BLAMEL4B, right). (D) Phylogenetic tree of 16S rRNA gene sequences of the methanogenic symbionts obtained in this study (bold), along with published sequences from cultured representatives as well as environmental sequences, and other published ciliate symbiont sequences. Symbiont sequences recovered in this study are colored by sample site (described in Figure S1, Table S1). Double dashes on branches indicate that branch was shorted to 50% of the original length. Bootstrapping was performed 1000 times and only values above 50 are shown. In (A)-(B), scale bar = 20 um. In (C), scale bar = 500 nm; MRO = mitochondrion related organelles; S = symbiont; black arrow = remnants of mitochondrial cristae; white arrow = mitochondrial double membrane.

**Figure 2.**
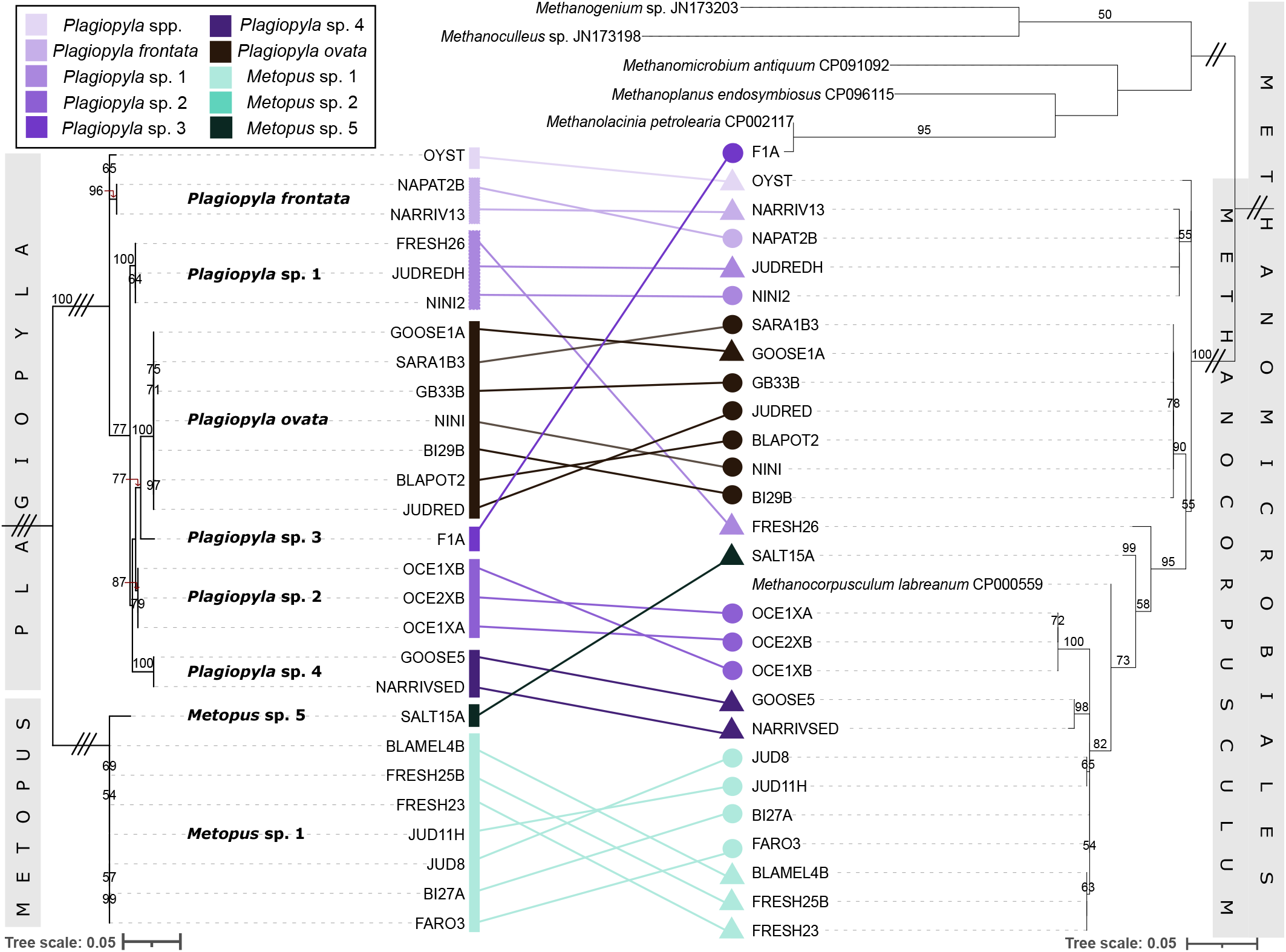
Paired host (left) and symbiont (right) phylogenetic trees. Host trees are based on the 18S rRNA gene sequences of the ciliates isolated in this study, and symbiont trees are based on the symbiont ITS region. Host and symbiont sequences are connected by lines, and lines and nodes are colored by host species group. Triple and double dashes on branches indicate that branches were shortened to 10% and 25% of the original length, respectively. Bootstrapping was performed 1000 times and only values above 50 are shown.

To provide higher resolution for distinguishing these symbionts (Figure 1), in addition to the 16S rRNA gene we also sequenced the symbiont 16S/23S internal transcribed spacer region (ITS). This is commonly used to detect strain-level differences among bacteria and archaea, including methanogens (18) (Figure 2). We found that *Plagiopyla* and *Metopus* host closely related yet distinct *Methanocorpusculum* endosymbionts, even when they co-occurred at the same location or in the same culture (e.g., FRESH25B and FRESH26; Figure 1, Figure 2). Additionally, symbionts cluster according to host species even when they originate from very distant locations or disparate habitats. For example, *Metopus* sp. 1 includes representatives isolated from the USA and in Portugal, and all its symbiont ITS sequences are identical (Figure 2). Symbionts of all *Plagiopyla ovata* populations are identical as well and include isolates from salt pond sediments in New England as well as mangrove sediments in Florida. Within host species, we sometimes see strain-level differentiation by geography: symbionts of *Metopus* sp. 1 isolates from sites 75 km apart formed site-specific clades despite hosts being virtually identical at the 18S rRNA gene level (FRESH and JUD, Figure 2). Even so, host-symbiont phylogenies are not congruent and, in some cases, *Metopus* and *Plagiopyla* symbionts are more related to each other than to symbionts from other species within their respective genera (Figure 2).

Here, we found that two, co-occurring, divergent marine ciliate lineages harbor very closely related *Methanocorpusculum* endosymbionts. These symbionts appear to be stable at the host species level, most likely through vertical transmission via synchronous division (10), but at higher taxonomic levels, there is evidence that symbiont swaps/replacements have occurred.

Mixed-mode transmission is common in cross-domain symbioses, especially those in aquatic environments (19). Particularly for marine symbioses, it has been found that this strategy can result in long-term evolutionary stability without extreme symbiont genome erosion via a combination of guaranteed transmission and symbiont replacement, while also allowing the host to associate with a locally adapted symbiont (20). Although it is ubiquitous in anoxia, this association is unique – the methanogenic symbionts of ciliates and other protists represent some of the few known intracellular archaea. Understanding the evolutionary dynamics of this association will provide insight into the complex mechanisms which shape the mixed-mode transmission strategies of cross-domain microbial symbioses. Future work using comparative and population genomics will be necessary to better help us understand how mixed-mode transmission shapes the ecology and evolution of these common yet complex partnerships.

## Supporting information

Tabl S4

## ACKNOWLEDGEMENTS

The authors would like to thank Daniel Méndez-Sánchez, Olga Čepičková, and Ivan Čepička, Sr. for kindly providing anaerobic ciliate cultures and micrographs, and to Veronica Berounsky, Virginia Edgcomb, Nicholas Cullo, and Vilém Helešic for help with obtaining samples from Rhode Island and Massachusetts sediments. This work was supported by a Simons Foundation Early Career Investigator in Marine Microbial Ecology, and Evolution Award to RAB as well as the United States National Science Foundation EPSCoR Track II Cooperative Agreement Award #1330406, and Czech Science Foundation grant 23-06004S awarded to IČ. The team acknowledges the Massachusetts Green High Performance Computing Center for computational resources and services. The TEM data was acquired at the RI Consortium for Nanoscience and Nanotechnology, a URI College of Engineering core facility partially funded by the National Science Foundation EPSCoR, Cooperative Agreement #OIA-1655221 and at the Viničná Microscopy Core Facility (VMCF of the Faculty of Science, Charles University), an institution supported by the MEYS CR (LM2023050 Czech-BioImaging). The S/TEM was purchased through an National Science Foundation Major Research Instrumentation grant #1919588.

## AUTHOR CONTRIBUTION

AS, JR, KP performed all cell culturing; AS JR performed all sample preparation and light microscopy; JR KP performed all electron microscopy imaging; KP prepared all of the electron microscopy samples; AS performed all phylogenetic analyses; AS JR RB IC conceptualized the study; RB IC acquired funding and supervised the co-authors; All co-authors discussed interpretation of the results; AS wrote the paper with input and revisions from all co-authors.

## DATA AVAILABILITY

Host 18S rRNA gene sequences for *Plagiopyla frontata* and *Plagiopyla ovata* are available on Genbank (accession numbers OP186382-OP186395, previously published). The remaining sequences (host 18S and symbiont 16S gene and ITS region sequences) are available on GenBank (accession numbers PP213518-PP213548, PP438670-PP438713, and PP215653-PP215679 and SUB14176396, respectively).

---

**Supplemental Figure 1.**
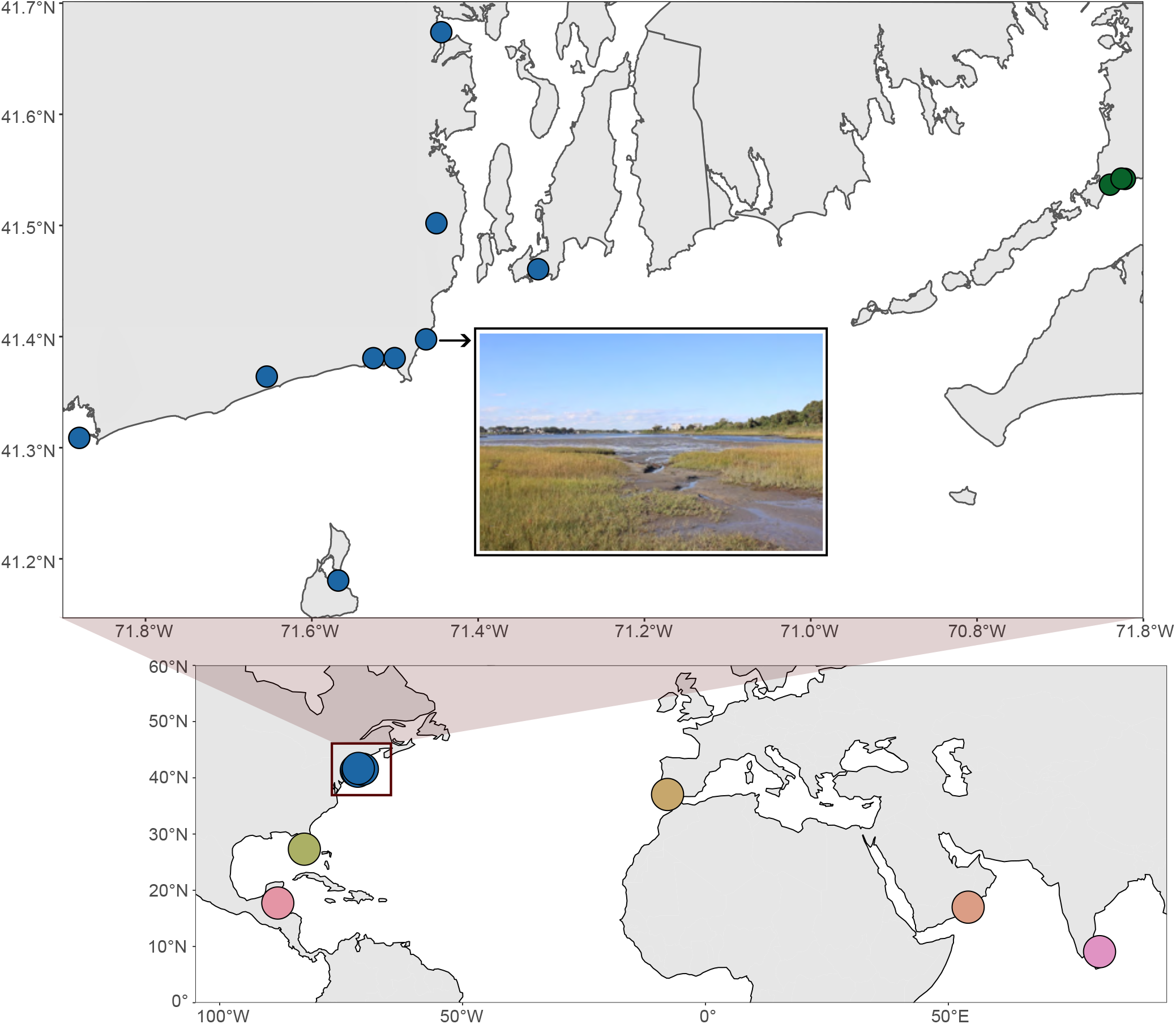
Map depicting localities and environments from which anaerobic ciliate populations from this study were obtained and cultured. Bottom panel shows samples isolated from locations globally, and top panel shows sample sites in Southern New England. Point Judith Pond, RI, USA sampling site is pictured in the top panel.

**Supplemental Figure 2.**
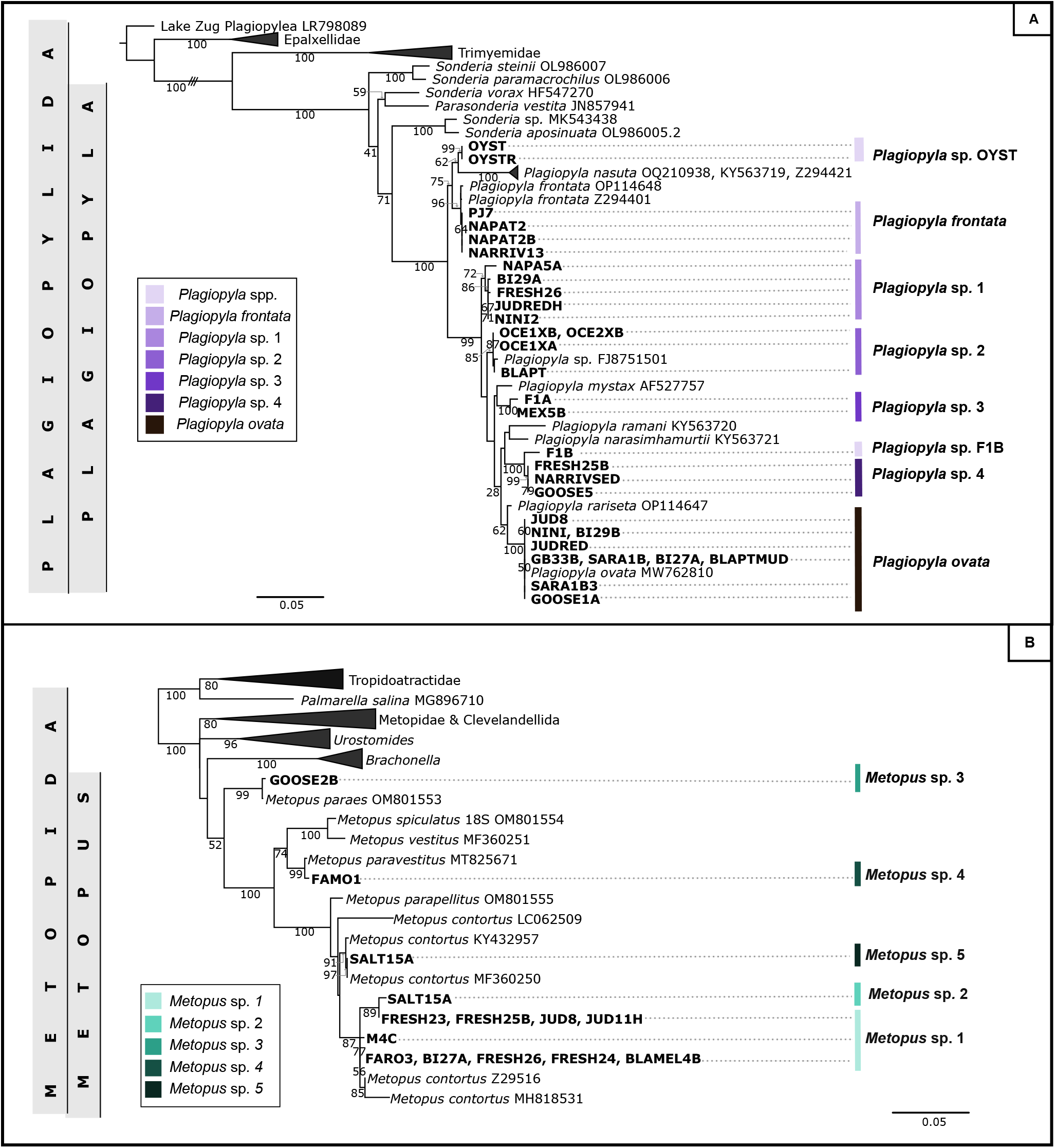
18S rRNA gene phylogenetic trees of host ciliate sequences isolated in this study (bold). (A) Phylogenetic tree of *Plagiopyla* sequences, along with representative sequences from Plagiopylea. (B) Phylogenetic tree of *Metopus* sequences, along with representative sequences from Metopida. Sequences are colored by species group. Triple dashes on branch indicates that branch wash shortened to 25% of original length. Bootstrapping was performed 1000 times and only values above 50 are shown.

## REFERENCES

1. Muñoz-Gómez SA. Energetics and evolution of anaerobic microbial eukaryotes. Nat Microbiol. 2023;8(2):197–203.

2. Husnik F, Tashyreva D, Boscaro V, George EE, Lukeš J, Keeling PJ. Bacterial and archaeal symbioses with protists. Curr Biol. 2021 Jul;31(13):R862–77.

3. Rotterová J, Edgcomb VP, Čepička I, Beinart R. Anaerobic Ciliates as a Model Group for Studying Symbioses in Oxygen-depleted Environments. J Eukaryot Microbiol. 2022 Sep;69(5):e12912.

4. Rotterová J, Salomaki E, Pánek T, Bourland W, Žihala D, Táborskỳ P, et al. Genomics of New Ciliate Lineages Provides Insight into the Evolution of Obligate Anaerobiosis. Curr Biol. 2020;

5. Fenchel T, Finlay BJ. The biology of free-living anaerobic ciliates. Eur J Protistol. 1991;26(3–4):201–15.

6. Worm P, Müller N, Plugge CM, Stams AJ, Schink B. Syntrophy in methanogenic degradation. In: (Endo) symbiotic methanogenic archaea. Springer; 2010. p. 143–73.

7. Lind AE, Lewis WH, Spang A, Guy L, Embley TM, Ettema TJ. Genomes of two archaeal endosymbionts show convergent adaptations to an intracellular lifestyle. ISME J. 2018;12(11):2655–67.

8. Fenchel T, Finlay BJ. Free-living protozoa with endosymbiotic methanogens. In: (Endo) symbiotic methanogenic archaea. Springer; 2010. p. 1–11.

9. Beinart RA, Rotterová J, Čepička I, Gast RJ, Edgcomb VP. The genome of an endosymbiotic methanogen is very similar to those of its free-living relatives. Environ Microbiol. 2018;20(7):2538–51.

10. Fenchel T, Finlay BJ. Synchronous Division of an Endosymbiotic Methanogenic Bacterium In the Anaerobic Ciliate Plagiopyla Frontata Kahl. J Protozool. 1991;38(1):22–8.

11. van Hoek AHAM, van Alen TA, Sprakel VSI, Leunissen JAM, Brigge T, Vogels GD, et al. Multiple Acquisition of Methanogenic Archaeal Symbionts by Anaerobic Ciliates. Mol Biol Evol. 2000 Feb 1;17(2):251–8.

12. Embley TM, Finlay BJ, Brown S. RNA sequence analysis shows that the symbionts in the ciliate Metopus contortus are polymorphs of a single methanogen species. FEMS Microbiol Lett. 1992;97(1–2):57–61.

13. Embley TM, Finlay BJ. The use of small subunit rRNA sequences to unravel the relationships between anaerobic ciliates and their methanogen endosymbionts. Microbiology. 1994 Feb 1;140(2):225–35.

14. Lynn D. The ciliated protozoa: characterization, classification, and guide to the literature. Springer Science & Business Media; 2008.

15. Li R, Zhuang W, Feng X, Al-Farraj SA, Schrecengost A, Rotterova J, et al. Molecular phylogeny and taxonomy of three anaerobic plagiopyleans (Alveolata: Ciliophora), retrieved from two geographically distant localities in Asia and North America. Zool J Linn Soc. 2023;zlad015.

16. Finlay BJ, Embley TM, Fenchel T. A new polymorphic methanogen, closely related to Methanocorpusculum parvum, living in stable symbiosis within the anaerobic ciliate Trimyema sp. Microbiology. 1993;139(2):371–8.

17. Volmer JG, Soo RM, Evans PN, Hoedt EC, Astorga Alsina AL, Woodcroft BJ, et al. Isolation and characterisation of novel Methanocorpusculum species indicates the genus is ancestrally host-associated. BMC Biol. 2023 Mar 22;21(1):59.

18. Ciesielski S, Bułkowska K, Dabrowska D, Kaczmarczyk D, Kowal P, Możejko J. Ribosomal Intergenic Spacer Analysis as a Tool for Monitoring Methanogenic Archaea Changes in an Anaerobic Digester. Curr Microbiol. 2013 Aug;67(2):240–8.

19. Russell SL. Transmission mode is associated with environment type and taxa across bacteria-eukaryote symbioses: a systematic review and meta-analysis. FEMS Microbiol Lett. 2019 Feb 1;366(3):fnz013.

20. Russell SL, Pepper-Tunick E, Svedberg J, Byrne A, Castillo JR, Vollmers C, et al. Horizontal transmission and recombination maintain forever young bacterial symbiont genomes. PLOS Genet. 2020 Aug 25;16(8):e1008935.

