## Supplemental Information for "Divergent marine anaerobic ciliates harbor closely related *Methanocorpusculum* endosymbionts"

**Supplementary information for "Divergent marine anaerobic ciliates harbor closely related *Methanocorpusculum* endosymbionts"**

Anna Schrecengost[1,a], Johana Rotterová[1,b], Kateřina Poláková[2], Ivan Čepička[2], Roxanne Beinart[1,a]

[1]Graduate School of Oceanography, University of Rhode Island, Narragansett, Rhode Island, USA

[2]Department of Zoology, Faculty of Science, Charles University, Viničná 7, 128 00 Prague 2, Czech Republic

[b]Current address: Department of Marine Sciences, University of Puerto Rico Mayagüez, Mayagüez, Puerto Rico, USA

**Contents**

**METHODS**

**SUPPLEMENTAL FIGURES AND TABLES**

**REFERENCES**

### METHODS

*Sample collection and culturing*

Sediment samples were collected by hand in 50-mL Falcon tubes from various locations in southern New England as well as Florida and various locations globally (Figure S2, Table S1). Each culture was established by inoculating 1-mL of sediment into 10-mL of Seawater Cereal Grass Media (ATCC #1525) for marine strains and into 10-mL of a mixture of seawater media and freshwater cereal grass media (ATCC #802) for brackish strains. Cultures were maintained by transferring the bottom 1-mL of culture into fresh media weekly/biweekly as described in (1).

*DNA extraction, PCR amplification, and sequencing of host and symbiont SSU rRNA genes*

For each strain, 30-100 ciliate cells were handpicked from culture with glass micropipettes, washed in sterile-filtered media, starved, and stored at -20°C in Zymo DNA/RNA shield (Zymo Research, Irvine, CA USA) as described in (2). Genomic DNA was extracted using the ZymoBIOMICS DNA microprep kit (Zymo Research, Irvine, CA USA) following the manufacturer's protocol. Host ciliate 18S rRNA genes from Plagiopylea strains were amplified using the general eukaryotic primers EukA (5'-AACCTGGTTGATCCTGCCAGT-3') and EukB (5'-TGATCCTGCAGGTTCACCTAC-3') and from Metopida strains using the Armophorea-specific primers ArmF1 (5'-GCGAYATRTCATTCAAGT-3') and ArmR4 (5'-GWGGTTWTCCACACAGTC-3')(3,4). Symbiont 16S rRNA genes were amplified using the methanogen-specific primers Met83F (5'-ACKGCTCAGTAACAC-3') and Met1340R (5'-CGGTGTGTGCAAGGAG-3') or Met86F (5'-GCTCAGTAACACGTGG-3') and 915R (5'-GTGCTCCCCCGCCAATTCCT-3') (5,6). The symbiont ITS region was amplified using the methanogen-specific primers 16S-RIS-M (5'-TGAAGCTGGAATSCGTAGTAATCGC-3') and

23S-RIS-M (5’-CTAAGATGTTTCAATYCVSNRSGTTCC-3’) (7). Products were purified with the enzymatic PCR cleanup reagent ExoSAP-IT Express (Applied Biosystems, Waltham, MA USA). Bi-directional Sanger sequencing was performed at the Rhode Island Genomics and Sequencing Center (Kingston, RI USA). Contigs were trimmed and assembled on Geneious Prime (https://www.geneious.com).

Partial 16S and 18S rRNA gene sequences from single cells of *Metopus* sp. 1 BLAMEL4B, BI27A, FRESH24, and FRESH26 and *Metopus* sp. 5 SALT15A were retrieved from single cell genomic data generated for another project. Briefly, the genomic data was generated from whole genome amplifications of single cells using the RepliG kit that were subsequently sent for 150-bp paired-end sequencing on a NovaSeq 6000 (Psomagen, MD, USA). Raw reads were trimmed with TRIMMOMATIC (8), cleaned, assembled with METASPADES (9), binned using MetaWrap (10), and finally retrieved from the respective *Methanocorpusculum* bins and host scaffolds via BLAST (11).

In vivo *microscopy, autofluorescence, and TEM*

The morphology of living cells was examined with a Nikon 80i epifluorescent microscope equipped with a digital single-lens reflex camera (Canon EOS Rebel T7i) and using differential interference contrast (DIC) and brightfield illumination. The presence of intracellular methanogens was confirmed *in-vivo* for each ciliate strain via autofluorescence of the methanogen-specific coenzyme F420 using with a UV filter (Excitation BP 395-440, Beam Splitter FT 460, Emission LP 470) according to Doddema and Vogels (1978). Transmission electron microscopy (TEM) was utilized for strains *Plagiopyla* sp. 1 FRESH26 and *Metopus* sp. 1 BLAMEL4B in order to localize the methanogenic endosymbionts inside of host cells.

Samples for TEM were processed as follows: samples were fixed in 2.5% glutaraldehyde for 4 hours at room temperature and postfixed in 1% osmium tetroxide for 1 hour on ice; after washing in phosphate-buffered saline and dehydration through a graded ethanol series, samples were transferred to acetone and embedded in EPON-Araldite resin. Ultrathin sections were cut on a Reichert-Jung Ultracut-E microtome (Reichert-Jung, Vienna, Austria) equipped with a diamond knife and contrasted with uranyl acetate and lead citrate. Sections were examined under a JEOL JEM-2100 transmission electron microscope equipped with an AMT XR401 sCMOS camera and under a JEOL-1011 transmission electron microscope equipped with a Veleta CCD camera.

*Phylogenetic analysis of host and symbiont 16S rRNA sequences*

For one of the symbiont phylogenetic trees, 16S rRNA gene sequences obtained from Sanger sequencing were aligned, along with reference and outgroup sequences obtained from GenBank, using the SILVA SINA aligner 1.2.11 (12). For the other symbiont tree, sequences from the symbiont ITS region obtained from Sanger sequencing were aligned, along with reference and outgroup sequences obtained from GenBank, using the MAFFT algorithm and the progressive method G-INS-1 (13). For the host trees, 18S rRNA gene sequences obtained from Sanger sequencing were aligned, along with reference and outgroup sequences obtained from GenBank, using the MAFFT algorithm and the progressive method L-INS-i. Reference sequences included 18S/16S rRNA gene sequences from previously published anaerobic ciliate/symbiont pairs. Alignments were manually trimmed to the primer regions using AliView (14). Phylogenetic trees were generated using a Maximum likelihood method in RaxML under the GTRGAMMAI model with 1000 bootstraps (15).

For the *Plagiopyla*, species-level lineages were determined based on average nucleotide identity using a threshold of 98.5% and on the formation well-supported clades in the 18S rRNA gene phylogeny (Figure S1, Tables S2 and S3). For the *Metopus*, species-level lineages were determined in the same way with the addition of morphological analysis (morphometrics on living and protargol stained specimens generated by William Bourland and Johana Rotterova for another project). Pairwise distance matrices for 18S and 16S rRNA genes were constructed using Clustal Omega in Geneious Prime (https://www.geneious.com) (16).

### SUPPLEMENTAL FIGURES

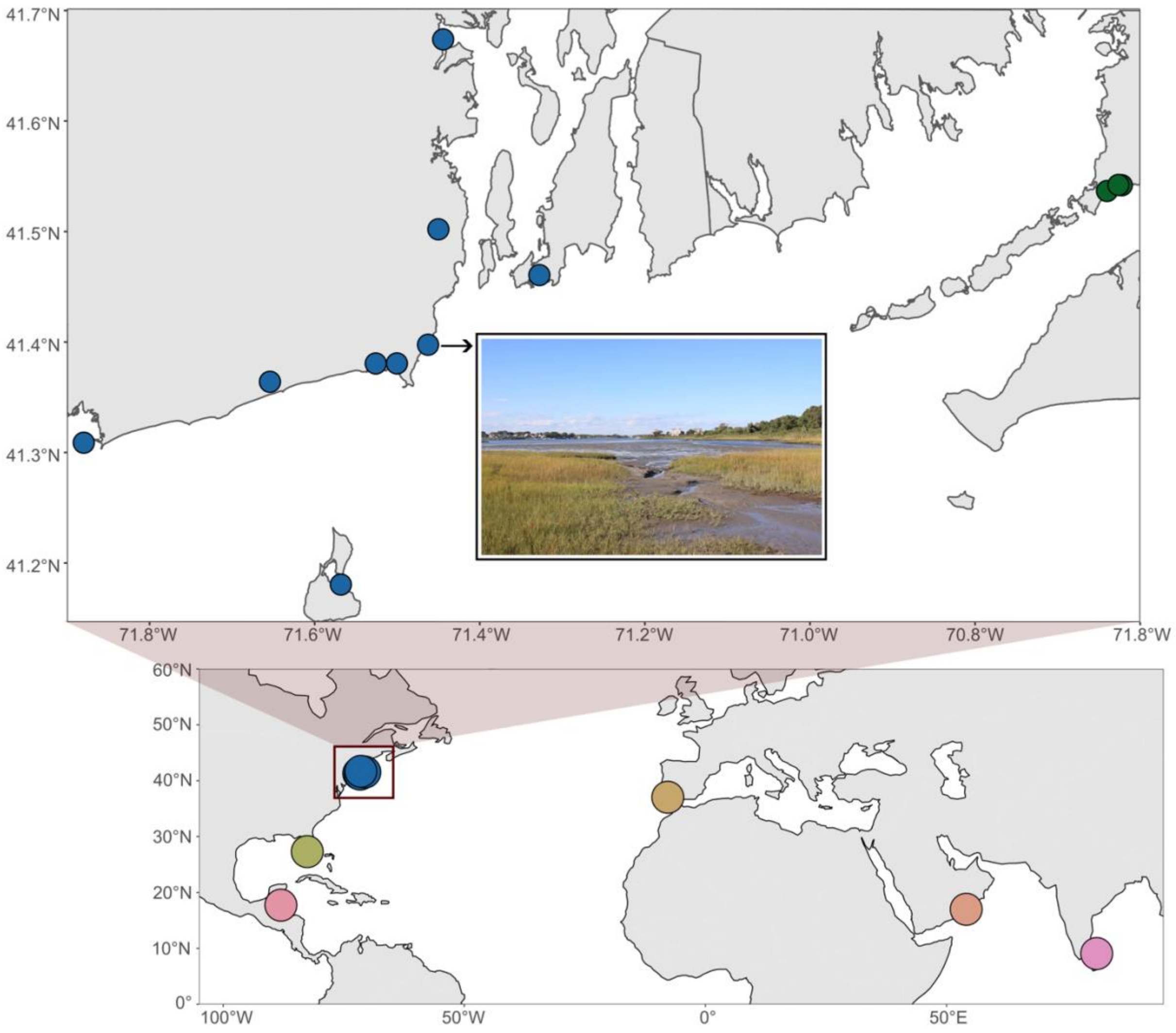


**Supplemental Figure 1.** Map depicting localities and environments from which anaerobic ciliate populations from this study were obtained and cultured. Bottom panel shows samples isolated from locations globally, and top panel shows sample sites in Southern New England. Point Judith Pond, RI, USA sampling site is pictured in the top panel.

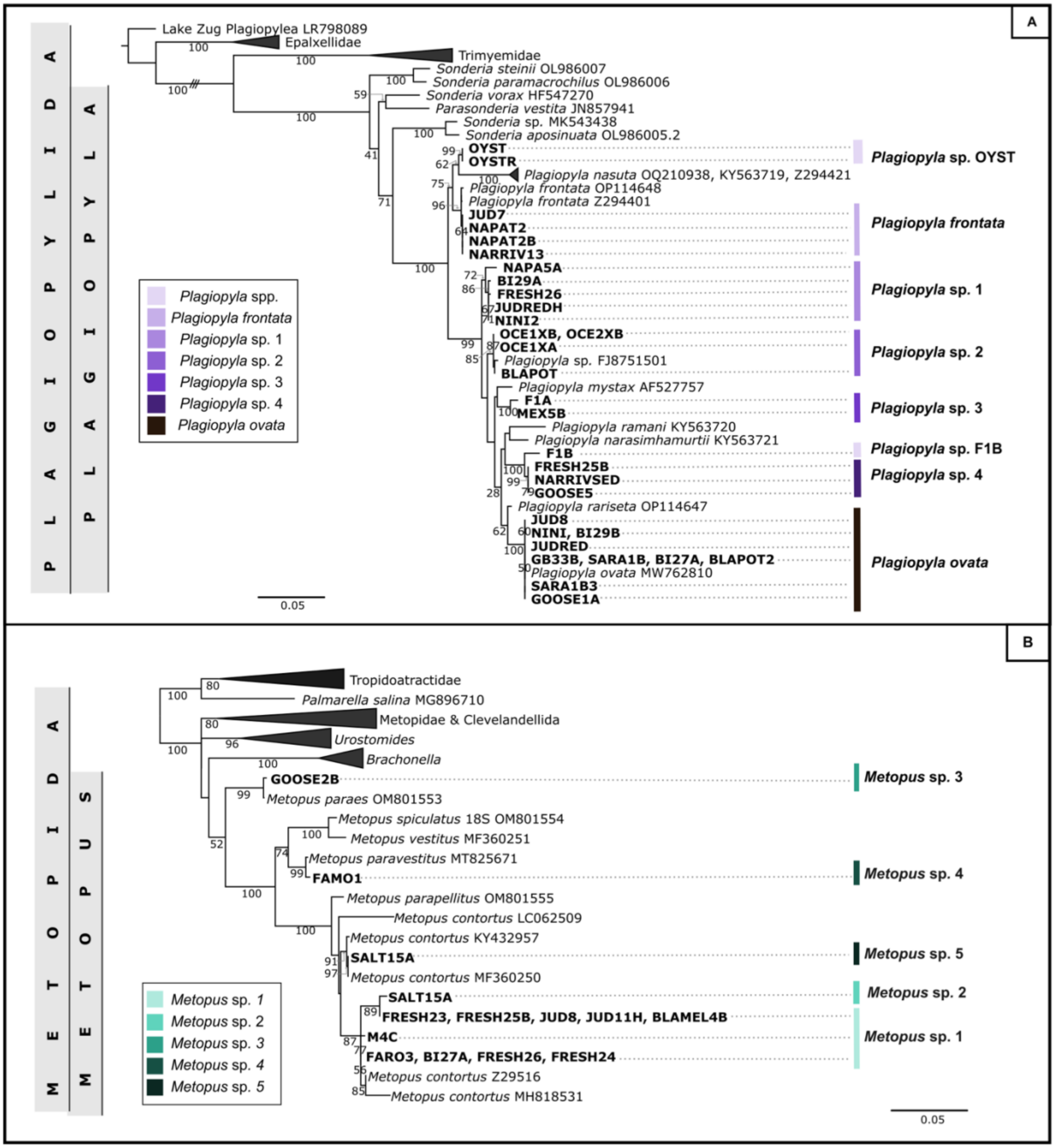


**Supplemental Figure 2.** 18S rRNA gene phylogenetic trees of host ciliate sequences isolated in this study (bold). (A) Phylogenetic tree of *Plagiopyla* sequences, along with representative sequences from Plagiopylea. (B) Phylogenetic tree of *Metopus* sequences, along with representative sequences from Metopida. Sequences are colored by species group. Triple dashes on branch indicates that branch wash shortened to 25% of original length.

### SUPPLEMENTAL TABLES

**Table S1.** List of anaerobic ciliate cultures established and analyzed in this study and associated metadata.

**Table S2.** Distance matrix of 18S rRNA gene sequences of *Plagiopyla* populations and closely related published sequences.

**Table S3.** Distance matrix of 18S rRNA gene sequences of *Metopus* populations and closely related published sequences.

**Table S4.** Distance matrix of 16S rRNA gene sequences of methanogenic symbiont populations and closely related published sequences.

### REFERENCES


1. Rotterova J, Čepička I. Cultivation protocol for anaerobic ciliates. 2019; Available from: https://www.academia.edu/download/87731276/cultivation-protocol-for-anaerobic-ciliates-85why7e.pdf
2. Beinart R, Rotterova J. Starvation & washing protocol for anaerobic ciliates. 2019 Nov 21; Available from: https://www.protocols.io/view/starvation-amp-washing-protocol-for-anaerobic-cili-868hzhw
3. Medlin L, Elwood HJ, Stickel S, Sogin ML. The characterization of enzymatically amplified eukaryotic 16S-like rRNA-coding regions. Gene. 1988 Nov 30;71(2):491–9.
4. Bourland W, Rotterova J, Čepička I. Redescription and molecular phylogeny of the type species for two main metopid genera, Metopus es (Müller, 1776) Lauterborn, 1916 and

Brachonella contorta (Levander, 1894) Jankowski, 1964 (Metopida, Ciliophora), based on broad geographic sampling. Eur J Protistol. 2017 Jun 1;59:133–54.

5. Wright ADG, Pimm C. Improved strategy for presumptive identification of methanogens using 16S riboprinting. J Microbiol Methods. 2003 Nov 1;55(2):337–49.
6. Tymensen LD, Beauchemin KA, McAllister TA. Structures of free-living and protozoa-associated methanogen communities in the bovine rumen differ according to comparative analysis of 16S rRNA and mcrA genes. Microbiology. 2012;158(7):1808–17.
7. Ciesielski S, Bułkowska K, Dabrowska D, Kaczmarczyk D, Kowal P, Możejko J. Ribosomal Intergenic Spacer Analysis as a Tool for Monitoring Methanogenic Archaea Changes in an Anaerobic Digester. Curr Microbiol. 2013 Aug;67(2):240–8.
8. Bolger AM, Lohse M, Usadel B. Trimmomatic: a flexible trimmer for Illumina sequence data. Bioinformatics. 2014 Aug 1;30(15):2114–20.
9. Nurk S, Meleshko D, Korobeynikov A, Pevzner PA. metaSPAdes: a new versatile metagenomic assembler. Genome Res. 2017 May;27(5):824–34.
10. Uritskiy GV, DiRuggiero J, Taylor J. MetaWRAP—a flexible pipeline for genome-resolved metagenomic data analysis. Microbiome. 2018 Dec;6(1):158.
11. Camacho C, Coulouris G, Avagyan V, Ma N, Papadopoulos J, Bealer K, et al. BLAST+: architecture and applications. BMC Bioinformatics. 2009 Dec;10(1):421.
12. Pruesse E, Peplies J, Glöckner FO. SINA: Accurate high-throughput multiple sequence alignment of ribosomal RNA genes. Bioinformatics. 2012 Jul 15;28(14):1823–9.
13. Katoh K, Rozewicki J, Yamada KD. MAFFT online service: multiple sequence alignment, interactive sequence choice and visualization. Brief Bioinform. 2019 Jul 19;20(4):1160–6.
14. Larsson A. AliView: a fast and lightweight alignment viewer and editor for large datasets.

Bioinformatics. 2014;30(22):3276–8.

15. Stamatakis A. RAxML version 8: a tool for phylogenetic analysis and post-analysis of large phylogenies. Bioinformatics. 2014;30(9):1312–3.

16. Sievers F, Wilm A, Dineen D, Gibson TJ, Karplus K, Li W, et al. Fast, scalable generation of high-quality protein multiple sequence alignments using Clustal Omega. Mol Syst Biol. 2011;7(1):539.
